# Towards AI-designed genomes using a variational autoencoder

**DOI:** 10.1101/2023.10.22.563484

**Authors:** Natasha K. Dudek, Doina Precup

**Affiliations:** School of Computer Science; McGill University; Montreal, QC, H3A 0G4; Canada; Mila - Québec Artificial Intelligence Institute; Montreal, QC, H2S 3H1; Canada

**Keywords:** Microbial genomics, machine learning, VAE, generative AI, bioinformatics

## Abstract

Genomes encode elaborate networks of genes whose products must seamlessly interact to support living organisms. Humans’ capacity to understand these biological systems is limited by their sheer size and complexity. In this work, we develop a proof of concept framework for training a machine learning algorithm to model bacterial genome composition. To achieve this, we create simplified representations of genomes in the form of binary vectors that indicate the encoded genes, henceforth referred to as genome vectors. A denoising variational autoencoder was trained to accept corrupted genome vectors, in which most genes had been masked, and reconstruct the original. The resulting model, DeepGenomeVector, effectively captures complex dependencies in genomic networks, as evaluated by both qualitative and quantitative metrics. An in-depth functional analysis of a generated genome vector shows that its encoded pathways are interconnected, near complete, and ecologically cohesive. On the test set, where the model’s ability to reconstruct uncorrupted genome vectors was evaluated, AUC and F1 scores of 0.98 and 0.83, respectively, support the model’s strong performance. This work showcases the power of machine learning approaches for synthetic biology and highlights the possibility that AI agents may one day be able to design genomes that animate carbon-based cells.

## INTRODUCTION

What is life? A fundamental goal of biology is to understand the principles that govern living systems. This involves mapping and understanding the function of molecular processes in cells, including how genes encode for these processes and how they interact with one another and their environment. The number and diversity of genes that have evolved in natural systems is vast, making this a formidable challenge. For example, it is estimated that 10^10^–10^13^ genes across 10^7^–10^8^ bacterial species have evolved since the dawn of life on Earth over four billion years ago [1,2].

Laboratory experiments can shed light on the nature of life, for example through the identification of genes that are essential for survival. Hutchison and colleagues employed transposon mutagenesis to remove non-essential genes from the *Mycoplasma mycoides* genome, which is amongst the most minimal that still encodes for cells capable of self-sufficient growth, reducing it from 901 genes to a mere 473 [3]. This reduction of life to its bare essentials was a breakthrough in our understanding of what constitutes life and was hailed as setting the stage for the design of life from scratch. Such efforts illustrate that while there is structure (i.e. sets of rules) that govern genome composition, elucidating said structure using laboratory techniques is labor intensive, time consuming, and costly; this particular experiment cost an estimated USD$40 million and required 20 people working for over a decade [4].

Machine learning (ML) excels at learning complex patterns in data. Generative ML models trained to produce synthetic genome designs could provide valuable insights into the genomic building blocks of life and the interactions between uncharacterized genes and metabolic networks. Further applications for such models include generating synthetic genomes for augmenting dataset size for developing ML models (akin to [5,6]), inferring metabolic potential of partially reconstructed genomes (similar to PICRUSt [7]), and computationally annotating genes in newly sequenced genomes. While the use of ML for *de novo* genome design remains largely unexplored, related tasks have benefited from the use of ML. Within synthetic biology, ML has shown promise for applications such as optimization of metabolite production levels [8–11], gene expression and metabolic flux [12–14], and fermentation [15–17]. Furthermore, generative ML approaches have been developed for the design of peptides (e.g., with anti-microbial, anti-cancer, or protein-protein interaction inhibition properties)[18–22], CRISPR guide RNA sequences [23], *de novo* antibodies [24–26], and synthetic promoter sequences [27].

Here, we asked the question: does generative ML have the potential to learn rules underlying genome composition? To explore this concept, we created a toy system of bacterial genomes and employed a denoising variational autoencoder (VAE) to model bacterial genome composition. First, we sourced 2,584 high-quality, complete bacterial genome assemblies from the KEGG database, and transformed each into a binary vectorial representation of the genes encoded in each genome, henceforth referred to as a genome vector. This simplification of a genome allowed us to perform an initial proof of concept study. To augment the dataset size while also preparing inputs for a reconstruction task, we created 100 heavily corrupted copies of each genome vector by masking bits representing genes. The VAE was then trained to reconstruct the original genome vector from the corrupted input, such that the output of the model is sets of genes that would be encoded for by viable bacterial cells. An in-depth metabolic analysis of a generated genome vector suggests that the model learns to generate largely complete, interconnected pathways, while an AUC score of 0.98 and an F1 score of 0.83 suggest that our model has high discriminative ability, precision, and recall. This work highlights the potential for ML algorithms to learn principles of genome composition.

## RESULTS

### Overview of DeepGenomeVector

In this work we propose and implement DeepGenomeVector, an ML approach to modeling the genetic composition of bacterial genomes. This model generates sets of genes that would be encoded for by viable bacterial cells. Gene sets are represented in the form of genome vectors, which are binary representations of all genes encoded by a complete genome. While this simplification was necessary to make the problem tractable for an initial study, it should be noted that genomes are far more complex than bags of genes; we did not model elements such as spatial organization of genes and non-coding regions (see Discussion).

Annotated bacterial genomes were sourced from a set of 2,584 complete, high-quality, annotated genomes from 46 major bacterial lineages (approximately phyla) from the KEGG genomes database [28–30] (Supplementary Figure 1A, B). To mitigate imbalances in the number of genomes across taxonomic groups, we thinned the genomic representation of overrepresented taxa (see Methods). Annotations in the form of KEGG Orthology (KO) identifiers were used to construct genome vectors consisting of 9,863 bits. While these KEGG annotations do not encompass the entirety of existing genes - an endeavor which is impossible due to the absence of known functions for many genes - they offer well-curated, consistent, and high-quality annotations for numerous metabolic and functional processes. Notably, genes in the KEGG database are organized into modules, which are functional units of gene sets. KEGG modules encode genetic functional units such as glycolysis, the ribosome, and methanogenesis. The fact that specific genes are parts of KEGG modules allowed us to more intuitively frame the task and observe aspects of the generated genome vectors that are especially important or valuable. The model, however, was not privy at any point to information about which genes are part of KEGG modules, nor to which genes belong to each KEGG module. Of the genes in our dataset, 11% (1,085 out of 9,863) were part of KEGG modules. Genome vectors in the dataset (training + test sets combined) encode a median of 1,885 genes, with a range of 528 - 4,536 genes per genome vector (Supplementary Figure 1C).

A major obstacle to using data-intensive ML methods to generate genome vectors is that the total number of complete, high-quality genome assemblies is limited. After splitting genomes into training (∼90%) and test (∼10%) datasets (see Methods) and converting them into genome vectors, we created 100 corrupted versions of every genome vector to augment dataset size.

Note that the choice of corruption process significantly impacts the results of a denoising algorithm by influencing the model’s ability to learn meaningful representations; more severe corruption typically encourages the learning of robust, abstract features, while less severe corruption tends to focus on finer, localized details [31]. In this study, the corruption process was designed to mimic potential real-world applications where an end user might want to generate synthetic genomes with particular functional properties, such as specific metabolic pathways or resistance mechanisms. Each corruption was achieved by retaining genes in 10 randomly selected KEGG modules encoded by an original genome vector, and masking all other genes. This encourages the model to learn both gene and functional unit co-occurrence patterns.

A denoising variational autoencoder (VAE) was selected from the family of generative machine learning algorithms for the task of modeling genome composition. In brief, VAEs learn to map their input to a latent space, sample from the latent space, and use that sample to generate a novel instance of the data. VAEs are trained to compress and decompress data, with latent space representations consisting of probabilistic distributions characterized by a mean and variance. The VAE code is then generated by sampling the probability distribution, and that sample is then decoded. Sampling of the distributions in the latent space is what confers the VAE its generative, rather than discriminative, properties [32]. A VAE was selected for this task primarily for its relative stability and ease of training compared to other generative models like GANs, which often present more complex training challenges. Importantly, their architecture allows for the direct comparison of inputs with outputs during training. This is significant as it enables the assessment of the similarity between an original, uncorrupted genome vector versus a generated genome vector, serving as a proxy for realism. A denoising architecture was selected to allow for the artificial augmentation of dataset size by creating corrupted versions of each genome vector and to encourage the model to learn more robust latent features.

The encoder and decoder of DeepGenomeVector each consist of three layers, with a latent representation size of 100 neurons (Figure 1), as this architecture was determined to yield optimal model performance during the training phase (see Methods). DeepGenomeVector was trained to optimize (i.e. minimize) the sum of binary cross-entropy (BCE) loss and

**Figure 1:**
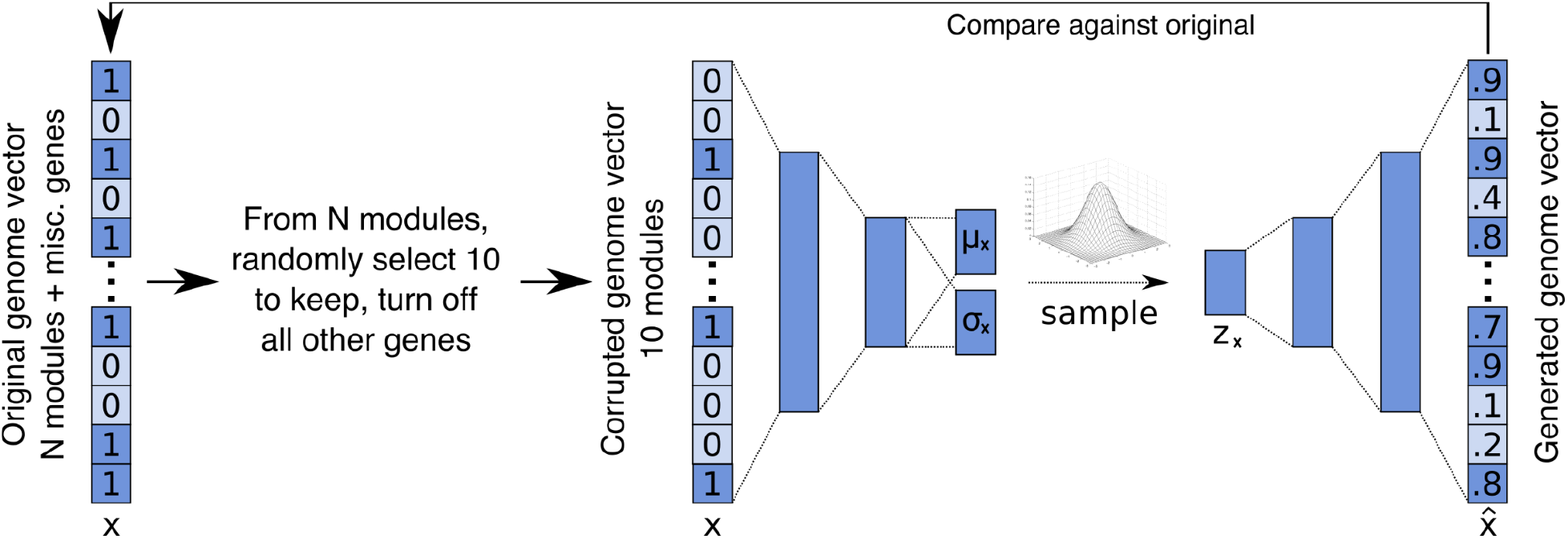
DeepGenomeVector’s architecture. Data corruption process and the DeepGenomeVector architecture. Genomes are represented as binary vectors (genome vectors) that denote which genes are encoded. To create input for the model, 100 corrupted versions of each genome vector are created by randomly selecting 10 functional modules it encodes, and creating a new corrupted genome vector that only encodes the genes from those 10 modules. The input to the model is a corrupted genome vector (x), where the presence (1) / absence (0) of genes is encoded. The model consists of three encoder layers (9,863, 500, 250, and 2 × 100 neurons) and three decoder layers (100, 250, 500, 9,863 neurons). The latent representation (z_x_) is sampled from the multivariate Gaussian distribution parameterized by μ_x_ and σ_x_. The output from the model is a probabilistic reconstruction of the corrupted input genome vector 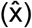. Per gene probabilities output by the model are binarized using a threshold of 0.5. The performance of the VAE is measured by comparing the generated genome vector against the original, uncorrupted one from which it was derived.

Kullback-Leibler divergence (KLD). Quantitative performance during development was evaluated using the F1 score, such that reconstructed genome vectors output by the VAE were compared against the original, uncorrupted genome vector from which the corrupted input to the model was derived.

Learning curves depicting loss and F1 score as evaluation metrics indicated a stable learning process (Figure 2A,B).

**Figure 2:**
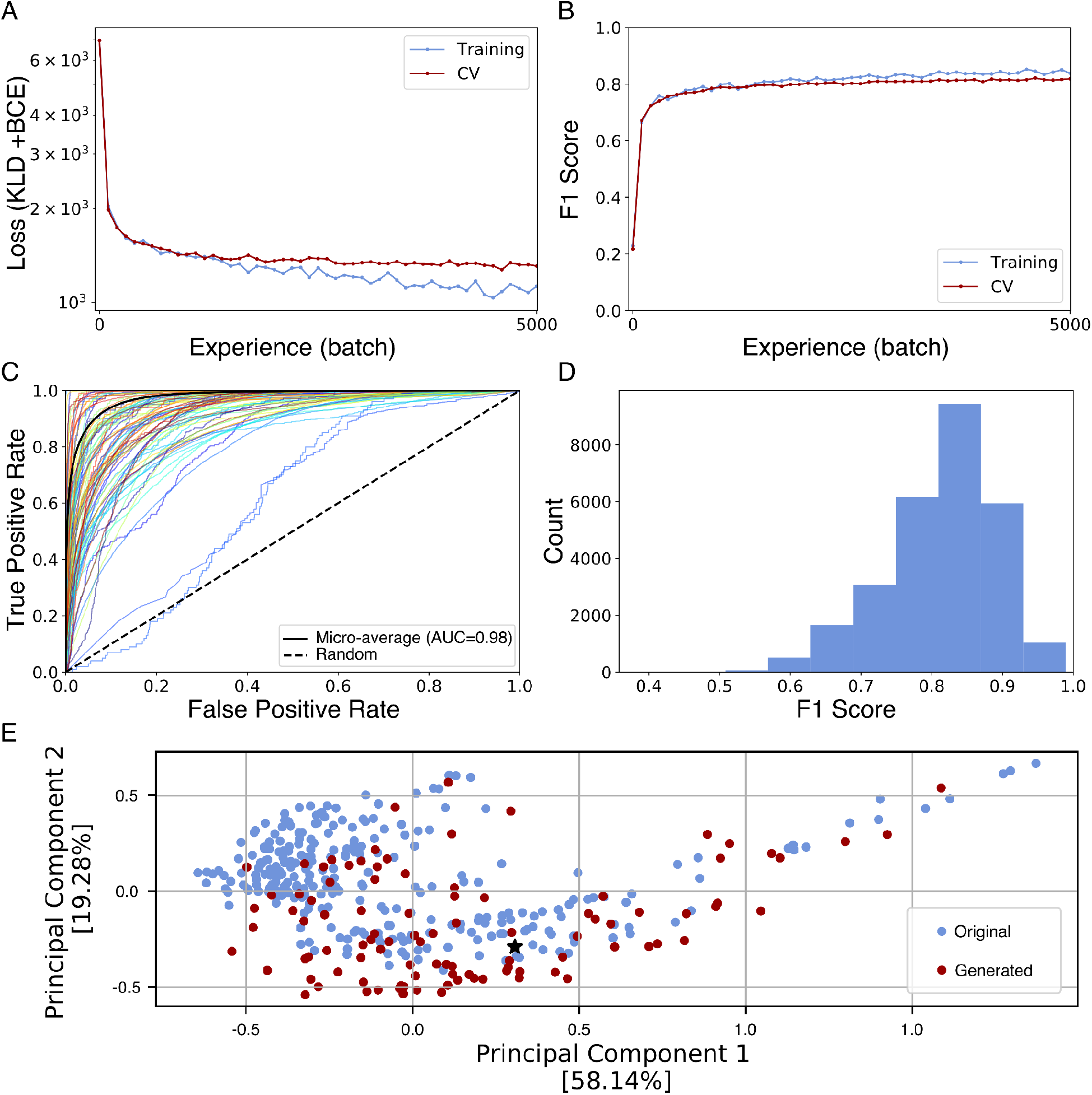
DeepGenomeVector learns and generates genome vectors that share substantial overlap in gene content with uncorrupted genome vectors. **(A)** Optimization learning curve, with batch loss (KLD + BCE) on the y-axis. Training was performed over 10 epochs. **(B)** Performance learning curve obtained using the test set, with F1 score on the y-axis. **(C)** Histogram of F1 scores obtained using the test set. **(D)** Receiver operating characteristic (ROC) curve obtained using the test set. Each coloured line represents one of 100 randomly selected genes (for clarity only a subset of the full 9,863 genes is visualized). The micro-average is plotted in solid black and corresponds to a micro-average AUC score of 0.98. **(E)** Principal component analysis (PCA) Jaccard similarity ordination of original (blue) versus generated (red) genome vectors. Jaccard similarity was calculated using the Hamming distance metric. Axes are scaled in proportion to the variance explained by each principal component. The generated genome vector selected for functional analysis (Figure 4) is denoted by a black star.

### Generated genome vectors overlap in genetic content with original genome vectors

We assessed the performance of DeepGenomeVector based on three criteria: realism, novelty, and diversity of generated genome vectors. Assessing the realism of a generated genome vector presents considerable challenges, as it involves determining the degree to which a given set of genes could theoretically support a living cell. As an initial approximation, we used the test set to quantitatively assess the degree to which reconstructed genome vectors were similar to the uncorrupted genome vectors from which they were derived. To gain insight into the model’s tradeoff between sensitivity and specificity per gene, we plotted an ROC curve (Figure 2C). A micro-average AUROC score of 0.98 suggests the DeepGenomeVector model has high discriminative ability, reflecting a high probability of correctly predicting the presence or absence of genes in a given genome vector. To further assess the predictive ability of the model, we computed F1 scores. The DeepGenomeVector model obtained F1 scores ranging from 0.99-0.39, with a median of 0.83 (Figure 2D). To contextualize the complexity of the task, we compared DeepGenomeVector’s performance against five baseline models that used heuristics to reconstruct genome vectors (Table 1). The baseline models were motivated and defined as follows. Baselines 1 and 2 were developed to assess the model’s performance relative to a naive strategy where outputs can be generated using limited to no meaningful data. Baseline 1 consisted of randomly turning on *n* genes in a corrupted genome vector from the test set, while baseline 2 consisted of randomly turning on the *n* genes with the highest probability of being on across the entire training set. The number of genes, *n*, was determined by randomly sampling from the number of genes present in original genome vectors across the training set. Baseline 3 was established as another reference point for the minimum expected performance, and consisted of using an untrained VAE to reconstruct corrupted genome vectors. Finally, baselines 4 and 5 were implemented to ensure that simply selecting a small or large number of genes was not a guaranteed strategy for success. Baseline 4 consisted of replacing corrupted data with genes from the most sparse genome vector in the training set (*Hoaglandella endobia*, Gammaproteobacteria), while baseline 5 consisted of replacing corrupted data with genes from the most dense genome vector in the training set (*Paraburkholderia caribensis*, Betaproteobacteria). The subpar performance of the five baselines on this task indicates that it is non-trivial, while the strong performance of DeepGenomeVector relative to these baselines suggests that it captures complex patterns within the data.

**Table 1:**
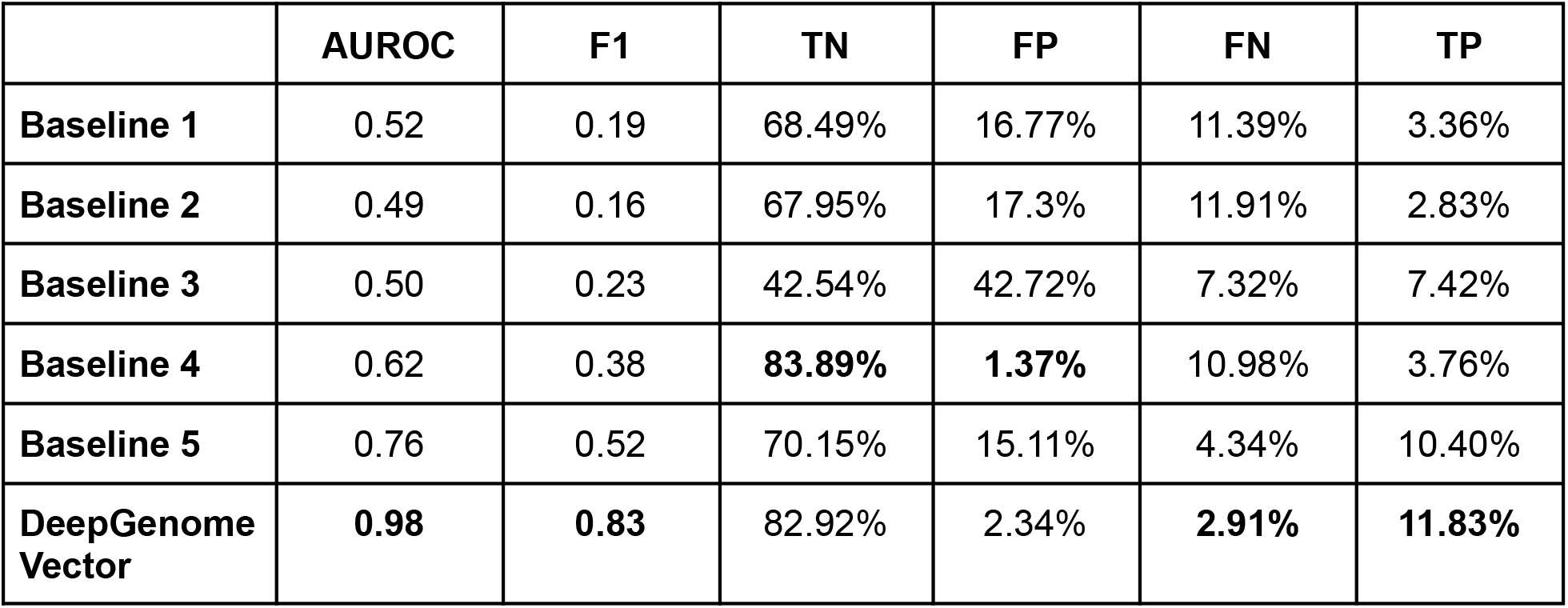
DeepGenomeVector’s outperforms baseline models on the test set. AUROC and F1 scores are micro-averages. True negatives (TN), false positives (FP), false negatives (FN), and true positives (TP) are the percentage of each, respectively, across all genome vectors in the test dataset. For baselines 1 & 2, let n = a randomly selected number of genes encoded by an original genome vector from the training set. Baseline 1 consisted of randomly turning on n bits in a corrupted genome vector from the test set. Baseline 2 consisted of randomly turning on the n bits with the highest probability of being on across the entire training set. Baseline 3 consisted of making predictions using an untrained VAE model. Baseline 4 consisted of replacing corrupted data with bits from the most sparse genome vector in the training set (*Hoaglandella endobia*, Gammaproteobacteria). Baseline 5 consisted of replacing corrupted data with bits from the most dense genome vector in the training set (*Paraburkholderia caribensis*, Betaproteobacteria).

|  | AUROC | F1 | TN | FP | FN | TP |
| --- | --- | --- | --- | --- | --- | --- |
| **Baseline 1** | 0.52 | 0.19 | 68.49% | 16.77% | 11.39% | 3.36% |
| **Baseline 2** | 0.49 | 0.16 | 67.95% | 17.3% | 11.91% | 2.83% |
| **Baseline 3** | 0.50 | 0.23 | 42.54% | 42.72% | 7.32% | 7.42% |
| **Baseline 4** | 0.62 | 0.38 | **83.89%** | **1.37%** | 10.98% | 3.76% |
| **Baseline 5** | 0.76 | 0.52 | 70.15% | 15.11% | 4.34% | 10.40% |
| **DeepGenome Vector** | **0.98** | **0.83** | 82.92% | 2.34% | **2.91%** | **11.83%** |

We next evaluated whether genome vectors with user-defined functional properties could be generated, and more specifically, whether the user-specified inputs to the model were reflected in the outputs it generated (Supplementary Figure 2). In 85% of cases, 90% or more of the original input genes were also present in the generated genome vector.

To gain further insight into whether generated genome vectors were qualitatively similar to original ones, we generated a set of 100 genome vectors for comparison against those from the test set. We calculated and plotted the Jaccard similarity between test genome vectors and a set of 100 generated genome vectors, using their Hamming distance, which is the number of bits that differ between two vectors (Figure 2E). While the distribution of generated genome vectors differs from that of original ones, there is significant overlap between them, and the absence of distinct clustering between generated and original genome vectors suggests that the generated genome vectors are qualitatively similar to the originals and fall within the natural range of variation.

Cumulatively, these results suggest that DeepGenomeVector excels at generating genome vectors that have substantial overlap in gene composition with original genome vectors.

### Generated genome vectors exhibit novelty and diversity

Preliminary evidence indicated that generated genome vectors exhibited similar gene content to original genome vectors from the test set. However, it remained to be determined whether DeepGenomeVector was a) closely mimicking the training set, or b) producing genuinely novel genome vectors rather than merely replicating the same outputs repeatedly. We first compared the similarity of generated genome vectors to the training data, using a distance metric. For the same set of 100 generated genome vectors as was used above, we calculated the Hamming distance from each generated genome vector to its closest neighbour in the training set (Figure 3A). For context, we repeated the same exercise comparing test set genome vectors against training set genome vectors. The distribution of minimum Hamming distances for generated genome vectors was greater than that for test set genome vectors. This suggests that generated genome vectors are more novel compared to those in the training set than are original genome vectors from the test set.

**Figure 3:**
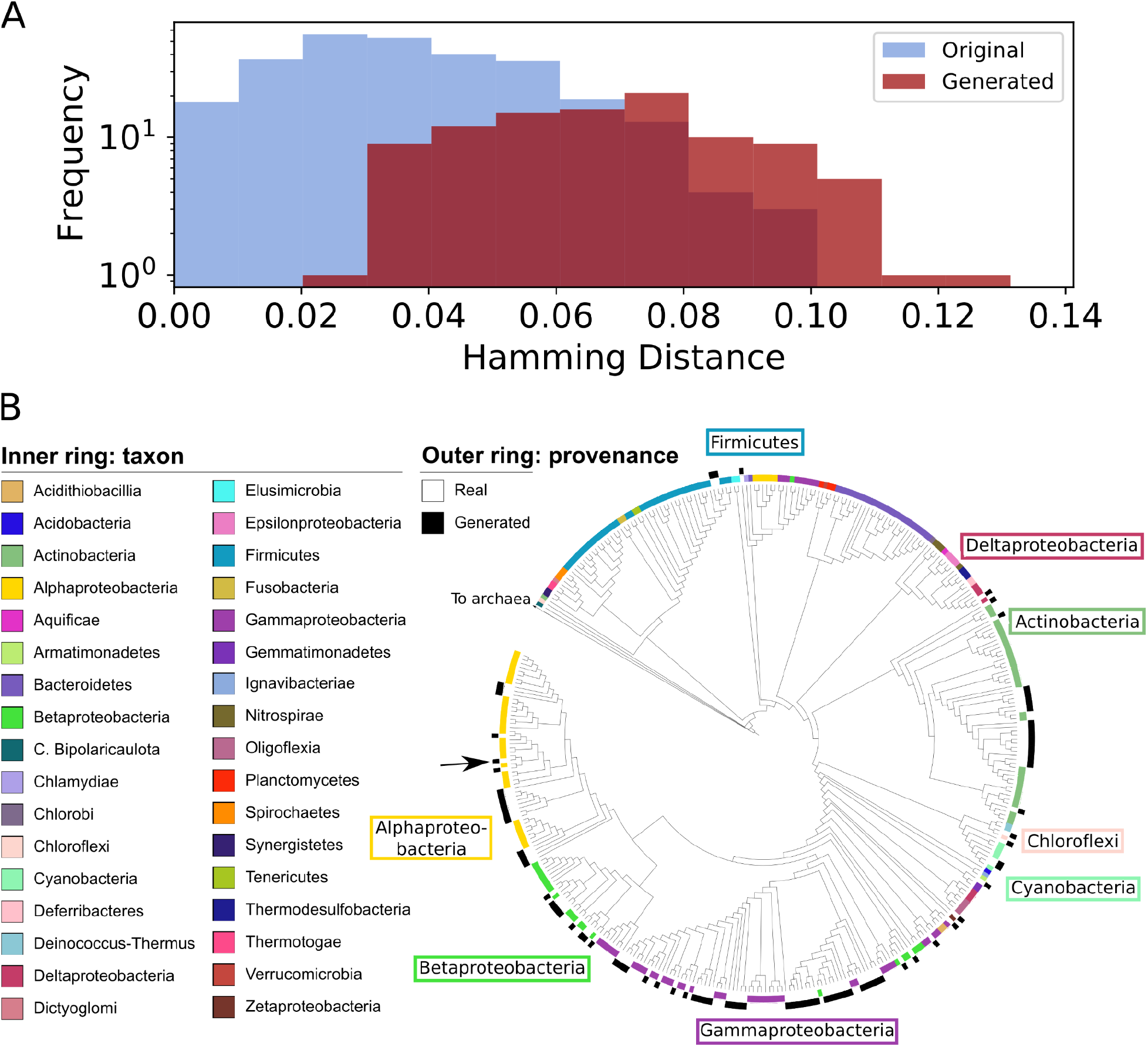
DeepGenomeVector’s generated genome vectors are novel and diverse. The 279 original genome vectors from the test set were compared against 100 generated genome vectors with randomly selected input modules. **(A)** Histogram showing the minimum Hamming distance of original genome vectors (blue) to the most similar training set genome vector versus generated genome vectors (red) to the most similar training set genome vector. **(B)** Parsimony-based dendrogram of original versus generated genome vectors. The dendrogram was generated from a character matrix where each gene’s presence/absence in a genome vector is a character. The inner ring around the dendrogram is a color-coded representation of genome vector phylum (original genome vectors only), while the outer ring indicates whether a genome vector was original (white) versus generated (black). Names of major lineages (approximately phyla) whose clades in the dendrogram neighbour or include generated genome vectors are indicated for reference. The generated genome vector selected for functional analysis (Figure 4) is denoted by a red arrow.

To assess the diversity of the genome vectors generated by DeepGenomeVector and contextualize their phylogenetic breadth, we incorporated them into a parsimony-based phylogenetic tree alongside the genome vectors from our test set. A dendrogram was built from a character matrix of gene presence/absence (Figure 3B), and shows that both original and generated genome vectors are widely distributed, with generated genome vectors often bounded within clades of original genome vectors (e.g.: Alphaproteobacteria). This supports that diverse genome vectors are being generated by DeepGenomeVector, representing lineages and lifestyles from across the bacterial tree of life (albeit skewed towards certain groups, likely reflective of the data imbalance in the training set and in nature itself).

### Metabolic reconstruction of a generated genome vector reveals largely complete and interconnected gene networks encoding for an ecologically plausible lifestyle

Finally, we investigated the extent to which metabolic pathways, cellular machinery, and other functional units were complete and properly connected, and whether lifestyles could be ecologically plausible. To gain insight into these questions, we performed an in-depth functional analysis of an example AI-generated genome vector, dubbed “g13” (generated #13), using techniques common to genome-resolved metagenomics [34–37] (Figure 4). This genome vector was selected quasi-randomly, with the sole criterion being that its position in the PCA ordination in Figure 2E (see black star) overlap with the distribution of original genome vectors.

**Figure 4:**
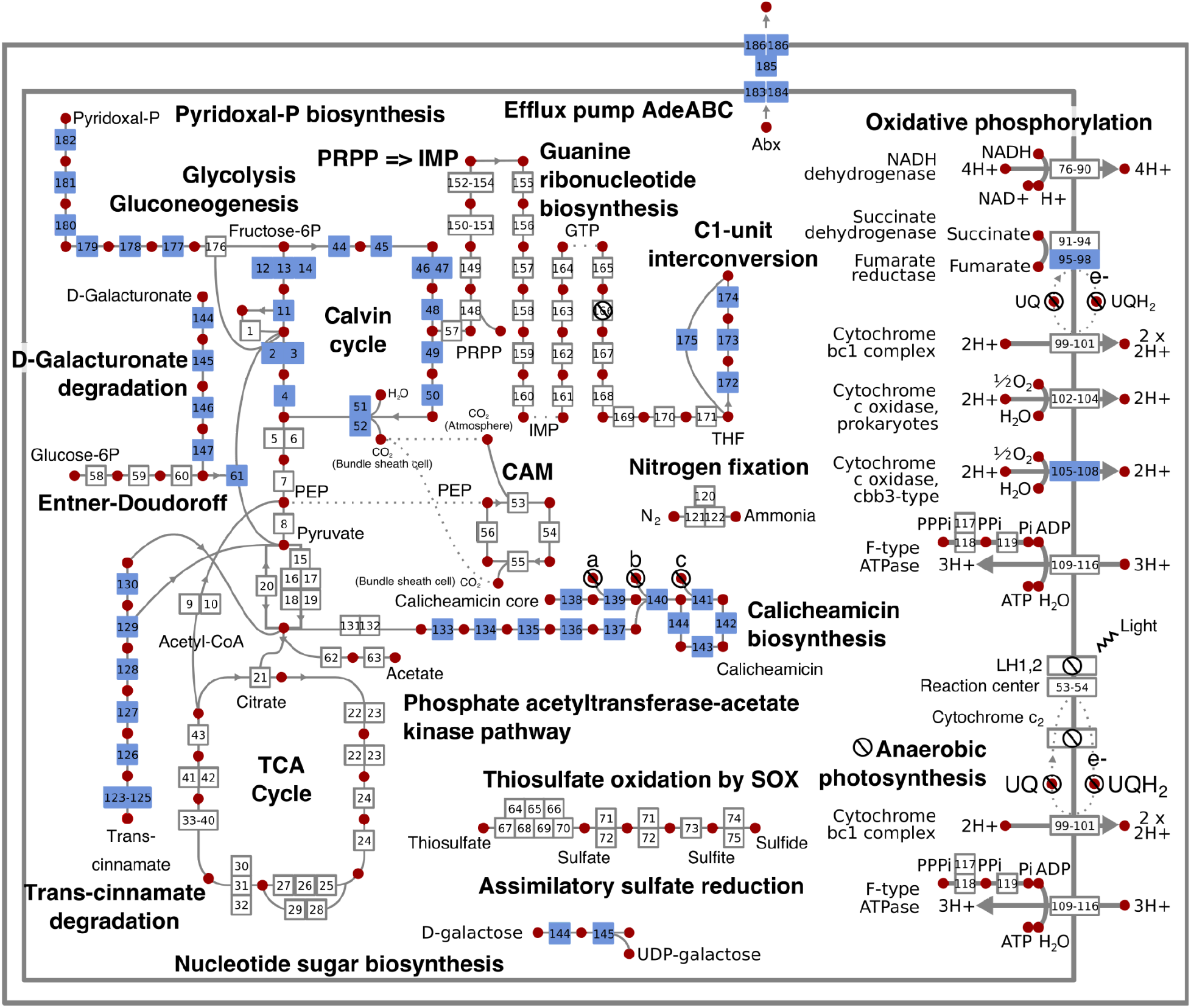
Cell diagram for a generated genome vector, depicting central carbohydrate metabolism, energy metabolism, and the 10 modules used as input. Boxes represent reactions catalyzed by genes encoded in the generated genome vector g13, while red dots represent key compounds involved in reactions. The 10 modules used as input to the model are colour-coded blue. Cross icons indicate genes that are not encoded or compounds whose synthesis is not encoded for. Metabolic pathways that are encoded by generated genome vector g13 but not directly related to central carbohydrate metabolism, energy metabolism, or an input module are not shown. Numbers in boxes allow for the identification of the reaction/gene via Supplementary Data 1.

First, we evaluated the extent to which KEGG modules were complete. Genome vector g13 encoded 2,015 genes, including 50/51 universal bacterial single-copy genes often used to assess bacterial genome completeness in metagenomics studies [38] (missing aspartyl tRNA synthetase). Core cellular machinery was near complete. For example, all genes required for an RNA degradosome were present, while 9/10 required for the RecFOR homologous recombination pathway were encoded. Of 30 KEGG modules encoded, only two were partial: guanine ribonucleotide biosynthesis (missing one gene) and anaerobic photosynthesis (missing light harvesting complexes 1 and 2, and cytochrome C2). This is striking when considering, for example, that all genes required for a complex process such as oxidative phosphorylation (n=41) were present. The genome vector lacks the ability to synthesize ubiquinol, however, which is a component of the electron transport chain.

We next evaluated the ecological plausibility of the lifestyle encoded by genome vector g13, by constructing a metabolic profile capturing its central carbohydrate metabolism, energy metabolism, and the 10 original input modules to the VAE (Figure 4; Supplementary Data 1). This is a small, meaningful subset of functional units that is manageable for human evaluation and aligns with one of the study’s objectives: designing genome vectors that encode specific, user-defined functional traits. Genome vector g13 encodes genes that would allow an organism to utilize compounds such as acetate and trans-cinnamate as carbon and energy sources. The Calvin cycle is also present, suggestive of a capacity for carbon fixation. The genome vector encodes glycolysis, the Entner-Doudoroff pathway, oxidative decarboxylation of pyruvate, the TCA cycle (forward direction), and oxidative phosphorylation, and would thus theoretically support aerobic respiration. The genetic potential to generate energy from inorganic compounds is present via both the assimilatory sulfate reduction and nitrogen fixation pathways. Anaerobic photosynthesis is partially encoded in g13. Encouragingly, modules were appropriately connected together, with inputs required for one pathway being produced by another. For example, the original input module C1-unit interconversion was connected to g13’s central carbohydrate and energy metabolism via the PRPP => IMP and guanine ribonucleotide biosynthesis pathway.The lifestyle encoded by genome vector g13 is reminiscent of a bacterium capable of both heterotrophic and lithoautotroph growth, and would be relatively metabolically versatile.

From this analysis, we conclude that AI generated genome vector 13 contains metabolic pathways and cellular machinery that are largely complete and properly connected together, and encodes ecologically cohesive metabolic pathways.

### Error analysis of DeepGenomeVector

To gain insight into model limitations, we examined biological and data-related factors that could potentially influence model performance. First, we queried whether the model performed better on genome vectors derived from certain phyla, or that encoded certain functional abilities. Using the test set, we calculated the median F1 scores of reconstructions derived from each phylum and encoding each KEGG module. Both phylum and KEGG module correlated with model performance (Kruskal-Wallis test, p = 0 and p = 1.01e^-154^, respectively), with Chlorobi-derived and photosynthesis-encoding genome vectors having the highest median F1 scores (Figure 5A,B). Notably, processes that are often acquired via horizontal gene transfer, such as drug resistance, tended to score poorly, as one might expect from modules not inherently tied to any given taxonomic group or lifestyle.

**Figure 5:**
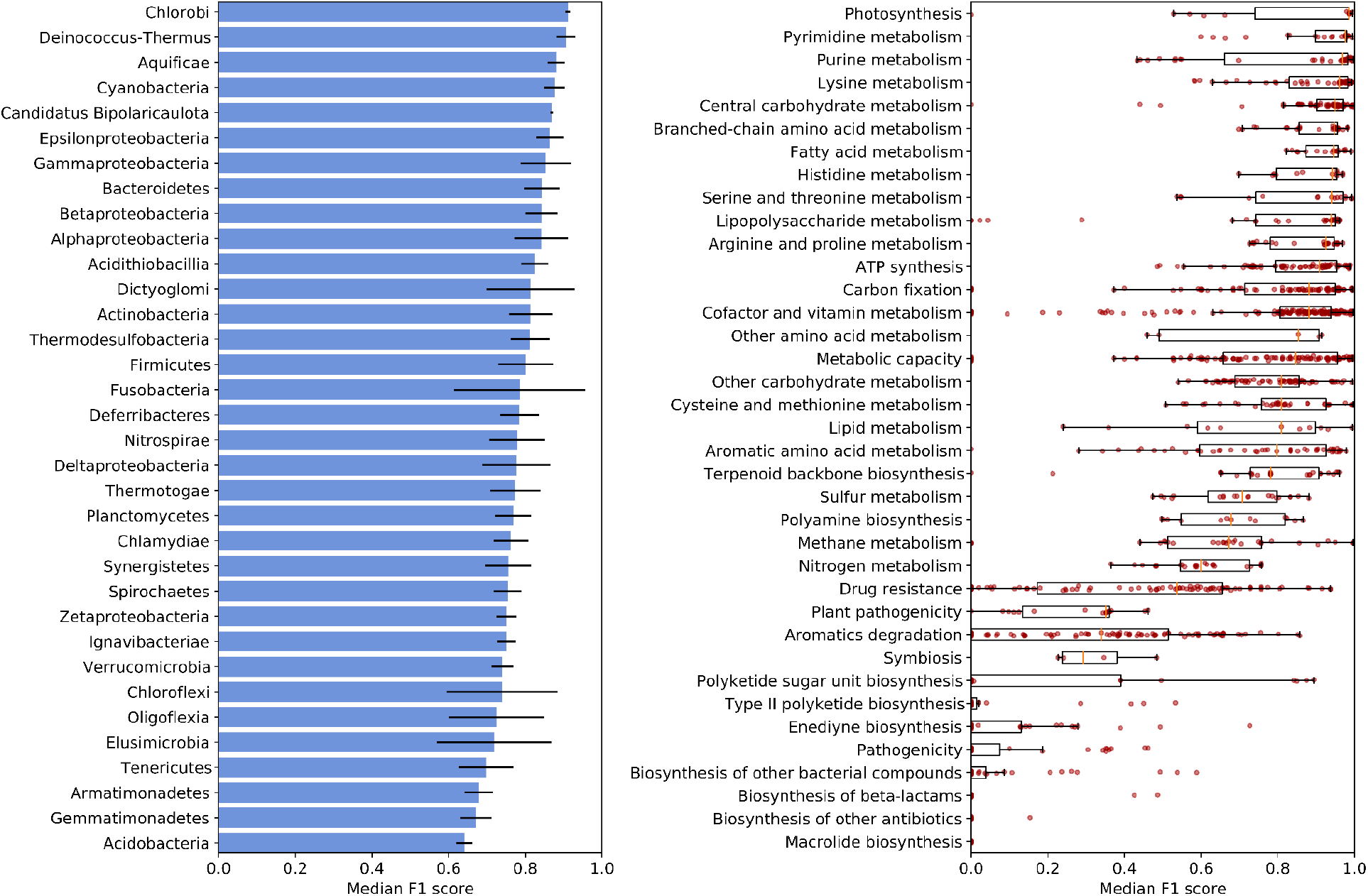
Biological covariates of DeepGenomeVector’s performance. **(A)** Model performance on genome vectors by taxonomic group (approximately phylum). Error bars represent median absolute deviation. **(B)** Model performance on genes by KEGG module. Individual data points represent modules. On the boxplots, the orange bars represent the median, boxes extend to the lower and upper quartile values, whiskers represent the range of the data, and data points beyond whiskers are outliers.

We next assessed whether DeepGenomeVector’s performance may be mechanistically correlated with properties of the data upon which was trained. We found that taxonomic composition of the training set at the genus level affected performance on the test set, but only when very few (none to one) same-genus representatives were present in the training set (ANOVA + Tukey test, p=5.98e-16). Similarly, genes that were less frequent in the training set were more likely to be mis-assigned during model evaluation (linear regression analysis, p=0.0). In both of these cases, it is likely that the composition of the training set reflects human sampling bias in genome coverage as well as unavoidable, naturally occuring discrepancies in the evolutionary diversity of certain taxonomic groups and the ubiquity of certain genes and lifestyles across the bacterial domain.

Finally, we examined the types of reconstruction errors to which DeepGenomeVector is susceptible, focusing on gene density and the inclusion of complete functional modules.

DeepGenomeVector appears to have learned a strategy whereby it uses more genes and complete genetic modules per genome vector than is typically found in an uncorrupted genome vector (Supplementary Figure 3), despite the fact that it was provided with no *a priori* information about which genes function together in any given module. Ecologically, this characteristic suggests that DeepGenomeVector generates genome vectors with increased metabolic versatility.

## DISCUSSION

In this study, we demonstrate the feasibility of using ML to model genetic principles that govern genome composition. When evaluating whether our model, DeepGenomeVector, was able to generate genome vectors similar in nature to those used as input seeds, AUC and F1 scores of 0.98 and 0.83, respectively, suggest high discriminative and predictive ability.

DeepGenomeVector substantially outperformed five heuristic baseline models, indicating its ability to capture and learn complex interdependencies in the data. Generated genomes appeared to be relatively realistic, novel, and diverse, as shown through distance based and phylogenetic assays. An in-depth functional analysis of a generated genome vector suggests that it is near-complete and encodes largely intact pathways that are interconnected, and would support an ecologically plausible lifestyle.

Although the model presented herein serves as a proof of concept for the modeling of genomic gene repertoires using machine learning, important limitations exist within this study. First, we acknowledge that there are substantial differences between a real genome and our simplified representation of one. Genome vectors do not represent many essential components of genomes, such as spatial organization of genes and non-coding regions (regulatory sequences, intergenic regions, etc). As such, DeepGenomeVector does not model the full complexity of genomic interactions. Furthermore, the features used for modeling only included genes annotated with a KO number from the KEGG dataset. This means our study excludes many genes that have evolved in nature but for which humans have not yet studied and characterized their function. Genes of unknown function can play essential roles in cellular biology [3]. Consequently, the DeepGenomeVector model omits a significant number of genes that exist in nature, including genes that are undoubtedly vital to cellular biology. Nonetheless, this proof of concept study shows that rules governing genome composition can be learned, even if those learned here cannot be complete due to limitations in current scientific knowledge. Notably, our approach may be extensible to aiding in the annotation genes of unknown function. DeepGenomeVector demonstrated an ability to learn the co-occurrence of genes within various metabolic pathways. If one were to cluster unannotated genes into groups of orthologs and model them, groups of interacting genes could theoretically be identified for functional studies. Another limitation of this work is that, to extend this work towards generating actual genomes rather than genome vectors, one would need to address how to translate binary vectors of 0s and 1s into gene sequences. The simplification of genome representations was necessary to make the problem manageable for an initial study; re-introducing complexity constitutes an area for future research.

An exciting progression of this work could be to impose constraints on models that mimic micro-evolutionary forces that shape bacterial genome architecture in nature [39]. For example, it has been theorized that resource specialization is driven by competition through natural selection [40,41]. At present, generated genome vectors have a tendency towards metabolically versatility, a trait which enables bacteria to function as generalists capable of adapting to diverse environmental conditions. Future efforts could focus on generating genome vectors that are metabolically sparse, which would align with the traits of specialists optimized for survival in narrow ecological niches. This could be accomplished by imposing sparsity constraints that reduce the metabolic diversity of output genome vectors.

In the future, it may be possible to improve model performance by employing GANs or even language-like models that generate sequences for the purposes of generating artificial genome vectors. A major barrier to doing so, however, is the limited amount of data available. Here, we circumvented this issue by creating corrupted versions of genome vectors for input to the model, which allowed us to train and evaluate the VAE. Critically, using a VAE allowed us to compare the original, uncorrupted genome vector against the genome vector generated from the corrupted input. This provided a rough evaluation of the quality of the generated product, albeit an imperfect one as many combinations of genes can be viable but may not be represented in the set available. Evaluation methods that are used to evaluate the realism of outputs of generative models in other fields, such as using the Frechet Inception Distance [33] for evaluating photo realism, require prohibitive amounts of data for this application. Methods for evaluating genome vector realism that are minimally data-hungry would broaden the scope of future approaches to genome vector design. Similarly, ever increasing dataset sizes will provide greater flexibility and breadth to ML applications in microbial genomics.

It is worth noting that the approach taken here can be adapted to new tasks by modifying the corruption process, thereby influencing the nature of the latent representations learned by the model. For example, a common task in genome-resolved metagenomics is to understand the functional profile of uncultured bacteria through the analysis of partial genome assemblies ([34–37]). If one were to extend the approach described here to assist in the prediction of a partial genome’s functional capabilities, it may be beneficial to implement a corruption process in which bits are randomly flipped. This could mimic the somewhat stochastic absence of various genome segments, and therefore genes, in a partially reconstructed genome. The change in corruption process would undoubtedly also affect the latent space representation learned by the model, as more severe or structured corruption (e.g., entire functional units) may encourage the model to represent abstract, high-level features in the latent space, whereas less severe or random corruption (e.g., randomly flipping bits) tends to promote the learning of specific, localized features [31]. Such considerations underscore the potential for numerous extensions to our approach.

DeepGenomeVector serves as concept validation that an ML algorithm can learn rules underlying genome composition. The development of such technologies may eventually lead to their use in synthetic biology, for example via their incorporation into synthetic genomic CAD software [42], or other applications such as dataset augmentation and computational genome annotation. From a more philosophical perspective, this work highlights the possibility that AI agents may one day be able to design viable carbon-based lifeforms.

## MATERIALS AND METHODS

### Genome selection, annotation, and representation

Genome annotations were acquired from the KEGG knowledge base [28–30]. The KEGG genomes database includes a set of 6,542 complete reference and representative genomes from NCBI that have been annotated using KEGG’s internal pipeline (accessed May 14, 2020, includes Eukaryotic, Archaeal, and Bacterial genomes). The KEGG bacterial genomes are strongly phylogenetically skewed; this reflects a natural imbalance in the known diversity of organism from different clades as well as their genomic representation in databases [43]. To mitigate the effect of this imbalance, we performed thinning to reduce the representation of over-represented taxa. For each genome, we used the taxid to recover the NCBI taxonomic identification. First, any genome not identified to the phylum level was discarded. Next we randomly selected only one genome to represent each species, and no more than five species to represent each genus. Per phylum we allowed for up to 50 genomes unclassified at the species level. Finally, we removed genomes with incomplete KEGG annotations, fewer than 500 annotated genes / genome (presumed symbionts with highly atypical lifestyles), and genomes encoding fewer than 10 KEGG modules (same rational re: symbionts). The final dataset consisted of 2,584 bacterial genomes spanning 46 major lineages (approximately phyla) and 9,863 features. For greater detail into the rationale behind these decisions, see the Supplementary Methods.

To represent genomes to an ML algorithm, we created binary vectors encoding whether, for each gene in the full dataset’s 9,863 genes, a given gene was encoded. These were termed genome vectors.

### Train-test split and corruption process

Genomes were split into training and test sets (∼90% and ∼10% of genomes, respectively) prior to undergoing conversion to vector format and corruption. As the goal of this study was to generate genome vectors representative of bacteria from across the bacterial domain, we included representatives from each bacterial phylum into each of the training and test sets where possible. This is important for ensuring the training and test sets are from the same distribution. A VAE trained only on generating genome vectors from phylum A will never generate high-quality genome vectors from a distantly related phylum B, much as a VAE trained to generate images of human faces (phylum Chordata) will not generate images of creatures from other eukaryotic phyla, such as worms (phylum Annelida), scorpions (phylum Arthropoda), or jellyfish (phylum Cnidaria). For major lineages (approximately phyla) represented in our dataset by ≥10 genomes, 88% were assigned to the training set and 12% to the test set. For those represented by 2 - 9 genomes, 50% were assigned to each of the training and test sets. Finally, for lineages represented by a single genome, that genome was assigned to the training set.

Corrupted genome vectors were used as input to the VAE. Corruptions were achieved by turning on the genes from 10 randomly selected KEGG modules that naturally co-occur within original genome vectors, and masking all other bits. This was performed 100X per genome vector in the training set and 100X per genome vector in the test set. The extent to which a genome vector was corrupted had a significant impact on the quality of generated genome vectors, as measured by the training set F1 score (Supplementary Table 1). We utilized the genes from 10 modules as input to minimize the number of genes a user would theoretically need to specify, while simultaneously mitigating the rapid decline in performance associated with utilizing genes from fewer than approximately 10 modules.

### Machine learning model

The DeepGenomeVector model was built in PyTorch v1.5.0 [44]. Hyperparameter tuning was implemented in Tune v0.8.2 [45] and performed using 10-fold cross-validation with AdamW as the optimizer [46,47]. Leaky ReLU activation functions [48] were applied to all layers except the output, for which a sigmoid activation was applied. Correspondingly, He initialization [49] was used for all layers except the last, for which Xavier initialization [50] was used. Sweeps over the following hyperparameters were conducted: number of layers (1, 2, 3, 4), batch size (32, 64, 128, 256), learning rate (log uniform selection between 1e^-4^ - 1e^-1^), and weight decay (log uniform selection between 1e^-5^ - 1e^-2^). While the number of layers had a strong impact on model performance, the effect of other hyperparameters was negligible. The final model used 3 layers for the encoder and decoder each, batch size 128, learning rate 0.001, and weight decay 0.1. At every iteration, loss was calculated by summing together binary cross-entropy (BCE) loss and Kullback-Leibler divergence (KLD). Training was performed over 10 epochs. When the model is used to generate a genome vector, the probabilities for each bit being on are converted to 0 or 1 using a 0.5 replacement threshold.

### Parsimony-based dendrogram

The 279 test genome vectors, 100 generated genome vectors, and an archaeal genome vector were used to create a parsimony-based dendrogram with the Wagner method [51,52].

This was achieved using the mix program in Phylip v3.695 [53]. The input order of species was randomized 10 times using seed=99 and using the archaeal genome vector as an outgroup root. All other settings were set to default. The dendrogram was visualized using iTOL v5.7 [54].

### Metabolic reconstruction of a generated genome vector

Identification of metabolic pathways was achieved using the KEGG Mapper reconstruct tool, available at https://www.genome.jp/kegg/mapper/reconstruct.html [55,56].

## CODE AVAILABILITY STATEMENT

Code for this project is available on Github at https://github.com/natasha-dudek/DeepGenomeVector.

## ACKNOWLEDGMENTS AND FUNDING SOURCES

This work was supported by NSERC Discovery and Canada-CIFAR AI Chair funding awarded to Doina Precup.

