## Supplementary Information for "Towards AI-designed genomes using a variational autoencoder"

**Supplemental information for**
**“Towards AI-designed genomes using a variational autoencoder”**

Dudek, N.K.[1,2], Precup, D.[1,2]

### AUTHOR AFFILIATIONS

1. School of Computer Science; McGill University; Montreal, QC, H3A 0G4; Canada
2. Mila - Québec Artificial Intelligence Institute; Montreal, QC, H2S 3H1; Canada

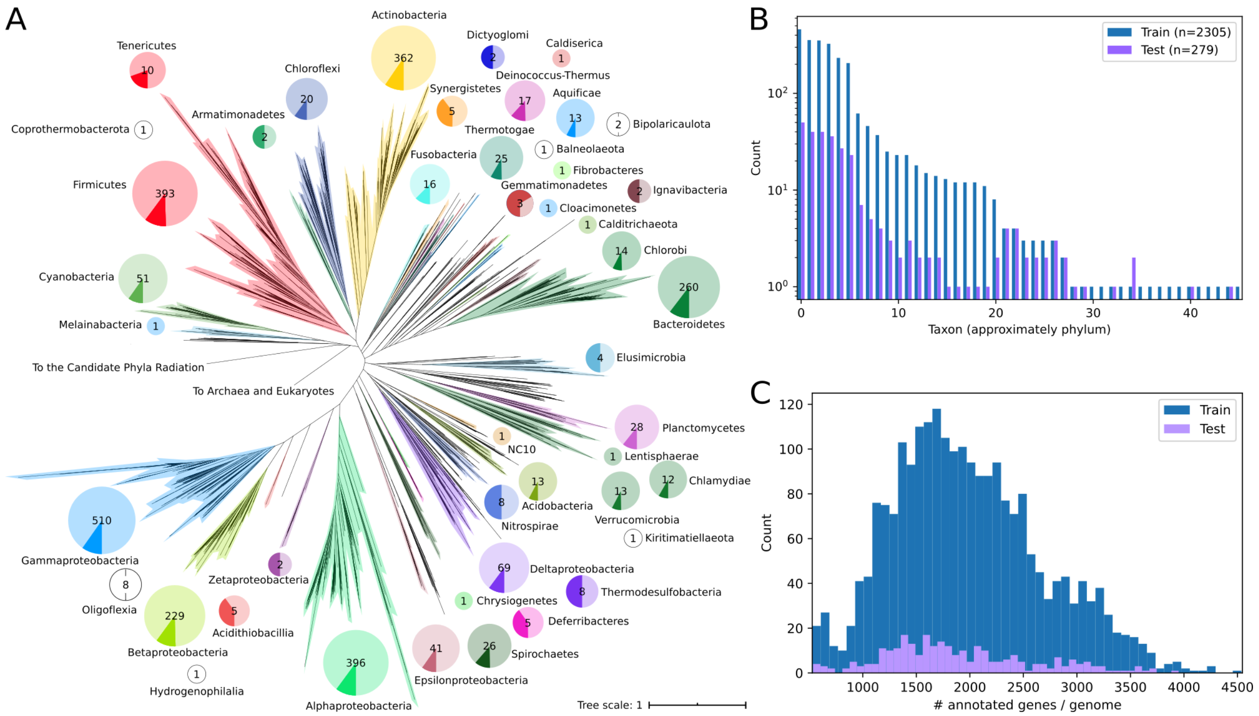


**Supplemental Figure 1: Overview of the genome dataset used to develop DeepGenomeVector. (A)** Phylogenetic distribution of genomes incorporated in the training and test sets. The bacterial tree of life is shown (modified from Hug *et al*., 2016), with the exclusion of the Candidate Phyla Radiation as no genomes from this supergroup were included. Major lineages (approximately phyla) from which genomes were included here are highlighted, with a colour-matching pie chart showing the portion of genomes from that lineage used in the training set (light) vs test set (dark). The total number of genomes from each lineage is indicated in the center of each pie chart. Major lineages not included in the Hug *et al*., 2016 tree are denoted by pie charts with no fill, and are placed in the proximity of the supergroup with which the given phylum affiliates. **(B)** The distribution of genomes per taxon (approximately phylum) in the training (blue) and test (purple) sets (see Methods). **(C)** Distribution of genes per genome. The number of genes annotated by KEGG per genome in the training (blue) and test (purple) sets are shown. Genomes with fewer than 500 annotated genes were not included in this study.

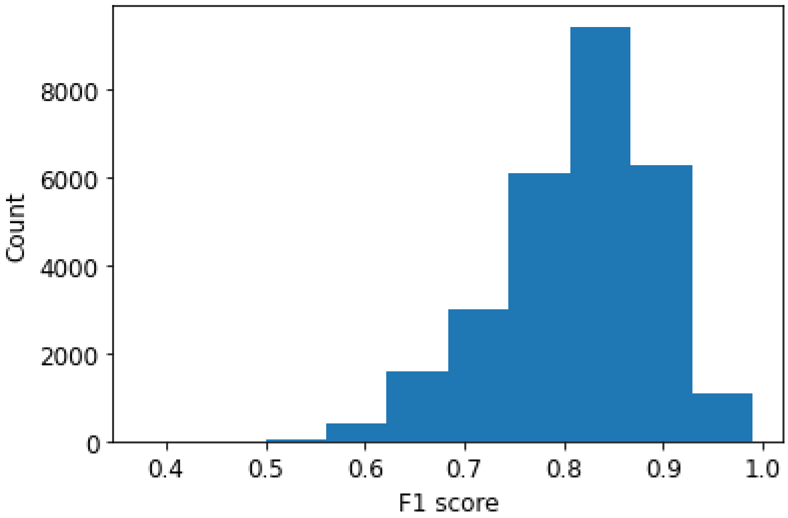


**Supplemental Figure 2: Distribution of DeepGenomeVector F1 scores across test set genome vectors.** F1 scores range from 0.38 - 0.99, with a median of 0.83.

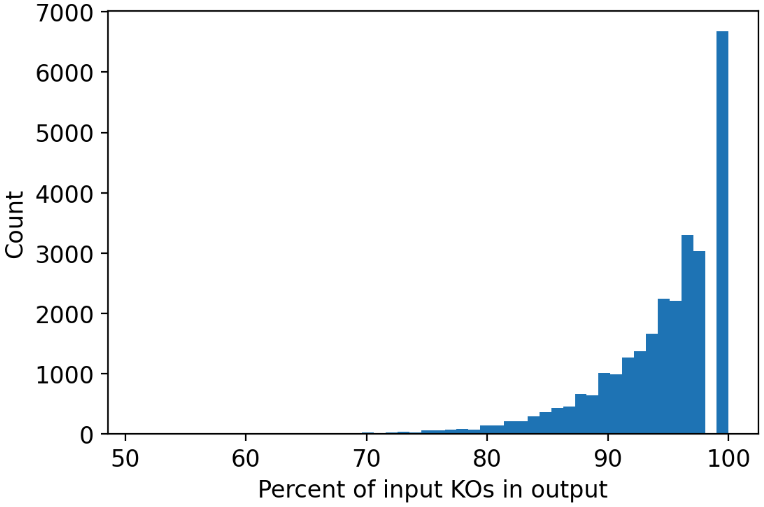


**Supplemental Figure 3: Frequency with which user-defined DeepGenomeVector inputs are present in the DeepGenomeVector output genome vector.** The percentage of input genes present in the output genome vector for a given genome is shown on the x-axis and occurrence counts are on the y-axis.

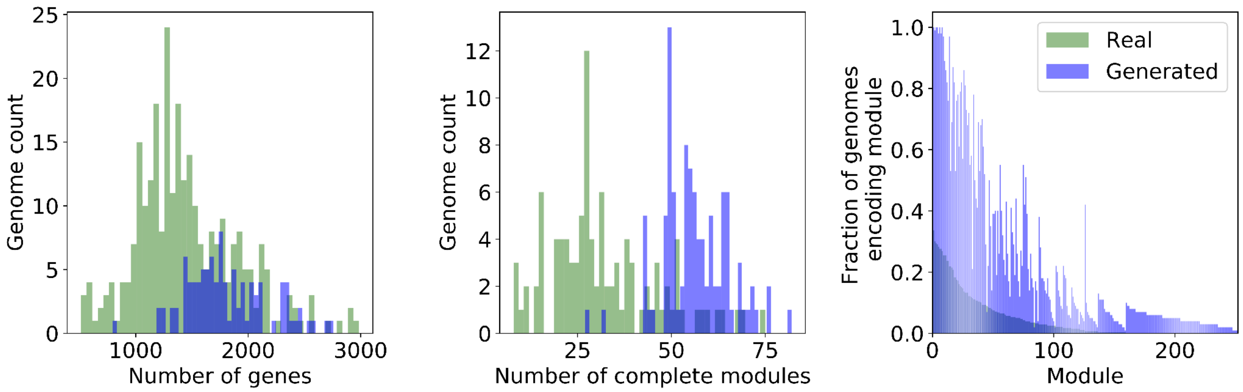


**Supplemental Figure 4: Distribution of gene and module counts for real vs. DeepGenomeVector generated genome vectors.** (**Left**) Distribution of the number of genes per genome. (**Center**) Distribution of the number of complete modules per genome. (**Right**) For each module, the fraction of genomes that encodes it.

### SUPPLEMENTAL TABLES

| # modules input | F1 score | # genomes | # features |
|---|---|---|---|
| 0 | 0.61 | 2332 | 9866 |
| 1 | 0.66 | 2330 | 9866 |
| 5 | 0.80 | 2325 | 9865 |
| 10 | 0.88 | 2305 | 9863 |
| 20 | 0.94 | 2200 | 9837 |
| 30 | 0.96 | 1955 | 9741 |
| 40 | 0.98 | 1580 | 9574 |

**Supplemental Table 1: Relationship between the number of modules retained during the corruption process and DeepGenomeVector F1 score.** Using the training dataset, the model was trained three times for each possible number of modules input. The best F1 score on any cross-validation fold across the three trials is reported. Because genomes with fewer than the threshold number of modules were discarded from the dataset during the corruption process, the number of genomes and features changes.

### SUPPLEMENTAL METHODS

#### Genome selection, annotation, and representation

For a machine learning algorithm to be able to learn what is encoded by a genome, it is essential that genomes used for training be complete, high-quality genomes that are consistently annotated such that homologues (i.e. the same gene in different genomes) are identifiable with a high degree of confidence. To acquire such annotated genomes, we turned to the KEGG knowledge base (1–3), which can also be used to link sequence information to high-level cellular functions. The KEGG genomes database includes a set of 6542 complete reference and representative genomes from NCBI that have been annotated using KEGG's internal pipeline (accessed May 14, 2020, includes Eukaryotic, Archaeal, and Bacterial genomes). Where possible, KEGG predicted proteins are assigned not only a name (e.g.: hexokinase) but also a KEGG Orthology (KO) number (e.g.: K00844). This is important because it allows for consistent identification and comparison of orthologous genes between genomes because each gene has a unique KO number, if not a unique name (e.g.: K00844 may be referred to as glucose ATP phosphotransferase, hexokinase type IV glucokinase, hexokinase (phosphorylating), etc). KEGG annotations are also of high quality as each KO number is assigned to protein sequence data in KEGG for which a protein function has been experimentally validated.

The KEGG bacterial genomes are strongly phylogenetically skewed. As such, we performed thinning to reduce the representation of over-represented taxa. For each genome, we used the taxid to recover the NCBI taxonomic identification. First, any genome not identified to the phylum level was discarded. Next we randomly selected only one genome to represent each species, and no more than five species to represent each genus. This worked for thinning well studied, well characterized lineages of bacteria, but there are many taxa for which this level of taxonomic classification is not available. For example, many candidate phyla do not robustly have defined classes, let alone genera or species. Thus per phylum we allowed for up to 50 genomes unclassified at the species level. Those lacking species classification were randomly selected to fill this quota. Thinning yielded 2,760 genomes, 250 of which were unclassified at the species level, and substantially helped even out the phylogenetic distribution of genomes, although it should be noted that the dataset is still phylogenetically skewed. This reflects that a) diversity is skewed across the tree of life (some lineages are more diverse than others), and b) genomic representation of diversity across the tree of life is skewed towards lineages that lend themselves to being cultured easily in the lab (Hug *et al.*, 2016). Finally, we removed genomes with incomplete KEGG annotations, fewer than 500 annotated genes / genome (presumed symbionts with highly atypical lifestyles compared to the “average” bacterium), and genomes encoding fewer than 10 KEGG modules (same rational re: symbionts). The final dataset consisted of 2,584 bacterial genomes spanning 46 major lineages (approximately phyla) and 9,863 features.
